# NeuroTri2-VISDOT: An open-access tool to harness the power of second trimester human single cell data to inform models of Mendelian neurodevelopmental disorders

**DOI:** 10.1101/2024.02.01.578438

**Authors:** Kelly J. Clark, Emily E. Lubin, Elizabeth M. Gonzalez, Annabel K. Sangree, Dana E. Layo-Carris, Emily L. Durham, Rebecca C. Ahrens-Nicklas, Tomoki T. Nomakuchi, Elizabeth J. Bhoj

## Abstract

Whole exome and genome sequencing, coupled with refined bioinformatic pipelines, have enabled improved diagnostic yields for individuals with Mendelian conditions and have led to the rapid identification of novel syndromes. For many Mendelian neurodevelopmental disorders (NDDs), there is a lack of pre-existing model systems for mechanistic work. Thus, it is critical for translational researchers to have an accessible phenotype- and genotype-informed approach for model system selection. Single-cell RNA sequencing data can be informative in such an approach, as it can indicate which cell types express a gene of interest at the highest levels across time. For Mendelian NDDs, such data for the developing human brain is especially useful. A valuable single-cell RNA sequencing dataset of the second trimester developing human brain was produced by Bhaduri et al in 2021, but access to these data can be limited by computing power and the learning curve of single-cell data analysis. To reduce these barriers for translational research on Mendelian NDDs, we have built the web-based tool, <u>Neuro</u>development in <u>Tri</u>mester <u>2</u> - <u>VI</u>sualization of <u>S</u>ingle cell <u>D</u>ata <u>O</u>nline <u>T</u>ool (NeuroTri2-VISDOT), for exploring this single-cell dataset, and we have employed it in several different settings to demonstrate its utility for the translational research community.

## Intro

Mendelian neurodevelopmental disorders (NDDs) are single-gene conditions that impact the development and function of the brain^1^. Forty percent of the more than 5,000 currently classified Mendelian disorders involve the brain or nervous system^2^, and over 1,500 genes have been implicated in these conditions^3^. Collectively, an estimated 9-18% of children are impacted by NDDs globally^3^, with approximately 3% of children diagnosed with a Mendelian NDD^1^. Every year more precise diagnoses are provided to families as about 300 new genes are identified as harboring causative variants underlying Mendelian NDDs^4^. Often, these novel NDDs fall into the classification of ultra-rare, impacting fewer than 1 in 50,000 individuals around the world^5^.

Putative disease candidate genes can be identified either through individual cases and small cohorts, or through the leveraging of large-scale publicly available datasets, such as gnomAD and GTEx^6–8^ (Figure 1A). Clinicians who provide care to these individuals can then connect with each other via resources like GeneMatcher and Matchmaker Exchange^9–11^. Through this approach, teams of clinicians and researchers can identify larger cohorts who share causative variants in the same genes and build more robust phenotypic profiles to inform care. Model systems are often not readily available to interrogate the pathogenic mechanism of these conditions and, even when they exist, these models can be financially prohibitive to obtain and space-prohibitive to maintain. Further, clinically accessible tissue (CATs) from affected individuals, like blood and skin, are useful for mechanistic studies but their availability is limited by research participation, logistical barriers and cost^12^ (Figure 1A). This lack of preexisting functional work and limited access to CATs for targeted assays can be a barrier to providing prognostic information to families.

**Figure 1.**
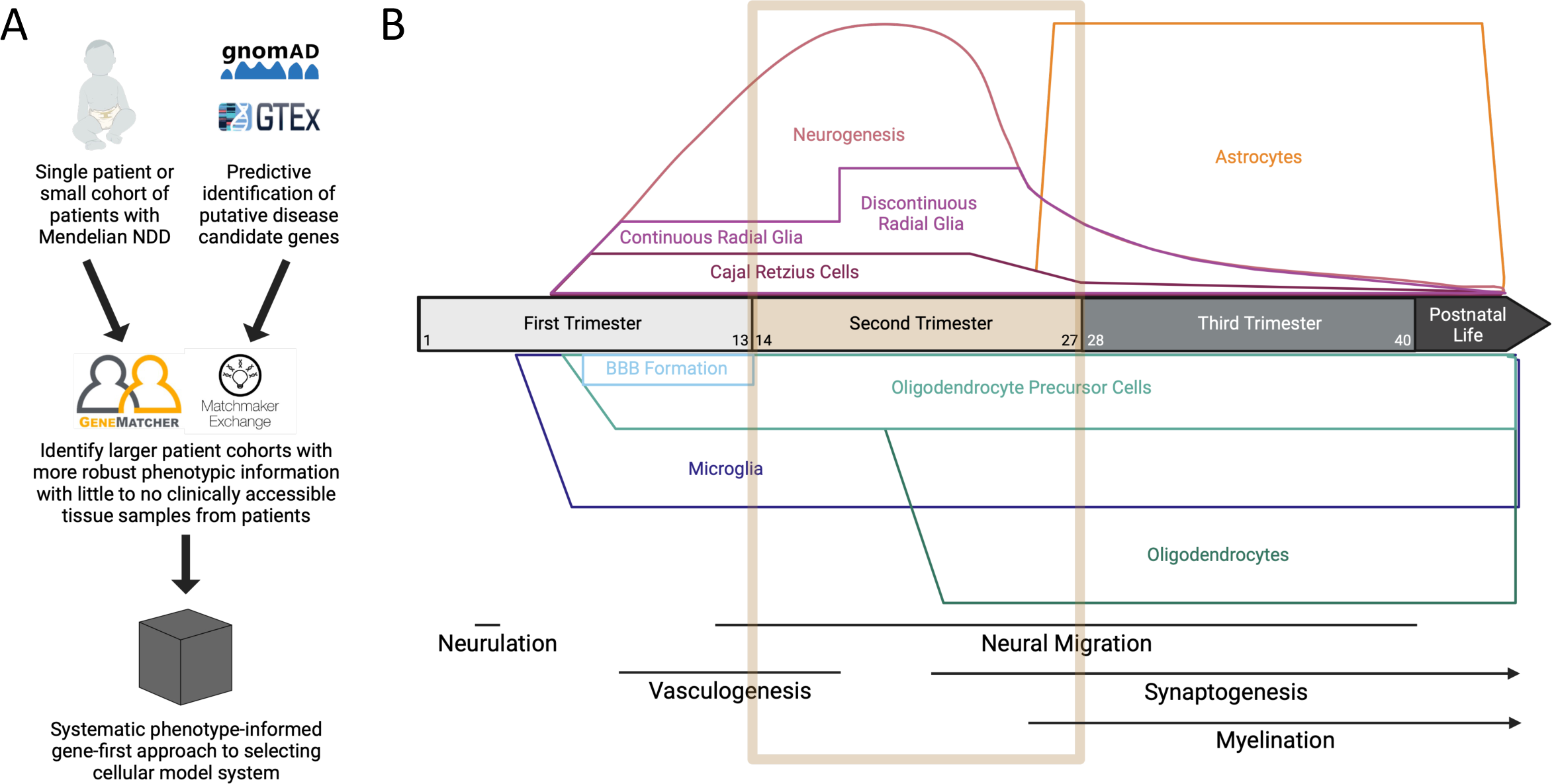
NeuroTri2-VISDOT fills key gap to inform selection of Mendelian NDD models. A) Proposed systematic approach to go from identification of putative Mendelian NDD gene to cohort identification to selection of the most useful model for downstream mechanistic work. Black box represents the gap filled by NeuroTri2-VISDOT. B) Timeline of human prenatal neurodevelopment including first- (gestational weeks (GW) 1-13; light gray), second- (GW 14-27; gold), and third-trimester (GW 28-40; gray), as well as the beginning of postnatal neurodevelopment (charcoal). All boxes/lines that extend past GW40 indicate continuance into postnatal life. Unfilled gold box highlights the second trimester of neurodevelopment, including the cell types present and the ongoing neurodevelopmental processes.

Thus, it is imperative for translational geneticists to have access to a systematic, phenotype-informed, genotype-first approach to select a model system. Human induced pluripotent stem cells (hiPSC) are powerful models, but it can be challenging to determine the most appropriate terminal lineage differentiation that recapitulates the pathogenesis of a particular ultra-rare Mendelian syndrome, especially if little to no functional work has been published for a gene of interest. Further, while model organisms like mice and zebrafish are incredibly powerful, especially for interrogating systemic effects of variants, obtaining and maintaining colonies of these organisms can be cost and time prohibitive. Additionally, there are fundamental differences in neurodevelopment between these organisms and humans that must be considered. Nonetheless, targeted exploration of specific cell types using *in vivo* systems can fill existing gaps in our current ability to differentiate hiPSCs to all the cell types that make up the human brain.

With the explosion of large-scale sequencing projects, an overwhelming number of datasets have been generated, and resources like the Chan Zuckerberg Initiative’s CELLxGENE platform help make these data more accessible^13^. Identifying the dataset that best aligns with a specific research question can still be a challenge. To specifically bridge the computational chasm for translational research questions motivated by individuals with ultra-rare Mendelian NDDs, we have built the tool <u>Neuro</u>development in <u>Tri</u>mester <u>2</u> - <u>VI</u>sualization of <u>S</u>ingle cell <u>D</u>ata <u>O</u>nline <u>T</u>ool (NeuroTri2-VISDOT). NeuroTri2-VISDOT references the powerful second trimester human fetal brain single-cell RNA-sequencing dataset generated by Bhaduri et al^14^, which is an important dataset for many reasons. First, these data cannot be widely generated, as access to fetal human brain tissue samples is limited. Additionally, generating and utilizing single cell RNA-sequencing data is a complex and computationally demanding process. The teams who produced these data have already overcome these two significant barriers to access. We sought to make these data more accessible so that the field of translational genetics can use them to responsibly inform selection of models of neurodevelopment, even without a robust computational background.

These second trimester data are an incredibly powerful window into human neurodevelopment (Figure 1B). The development of the human central nervous system (CNS) begins after gestational week 3 (GW3), following the closure of the neural tube, a process called neurulation (Figure 1B)^15^. During the first trimester, neural immune cells like native cells and microglia colonize the CNS and begin to proliferate prior to the formation of the blood brain barrier^16–19^ (Figure 1B, Table 1). Neurogenesis and gliogenesis also begin in the first trimester but, at this stage, a limited number of cell types are identifiable based on transcriptomic profiling, including radial glia (RG), neural progenitor cells (NPCs), and neurons^20^. That stands in contrast to what can be delineated in the second trimester, where transient prenatal cell types, such as Cajal-Retzius cells, as well as many of the CNS cell populations that persist into post-natal life are captured (Figure 1B, gold box; Table 1). Processes essential to neurodevelopment also occur during the second trimester. For instance, RG undergo the human-specific process of becoming a physically discontinuous scaffold between GW16.5-17, which is not captured in commonly employed model systems of neurodevelopment, such as mice or zebrafish^21,22^.

**Table 1.**
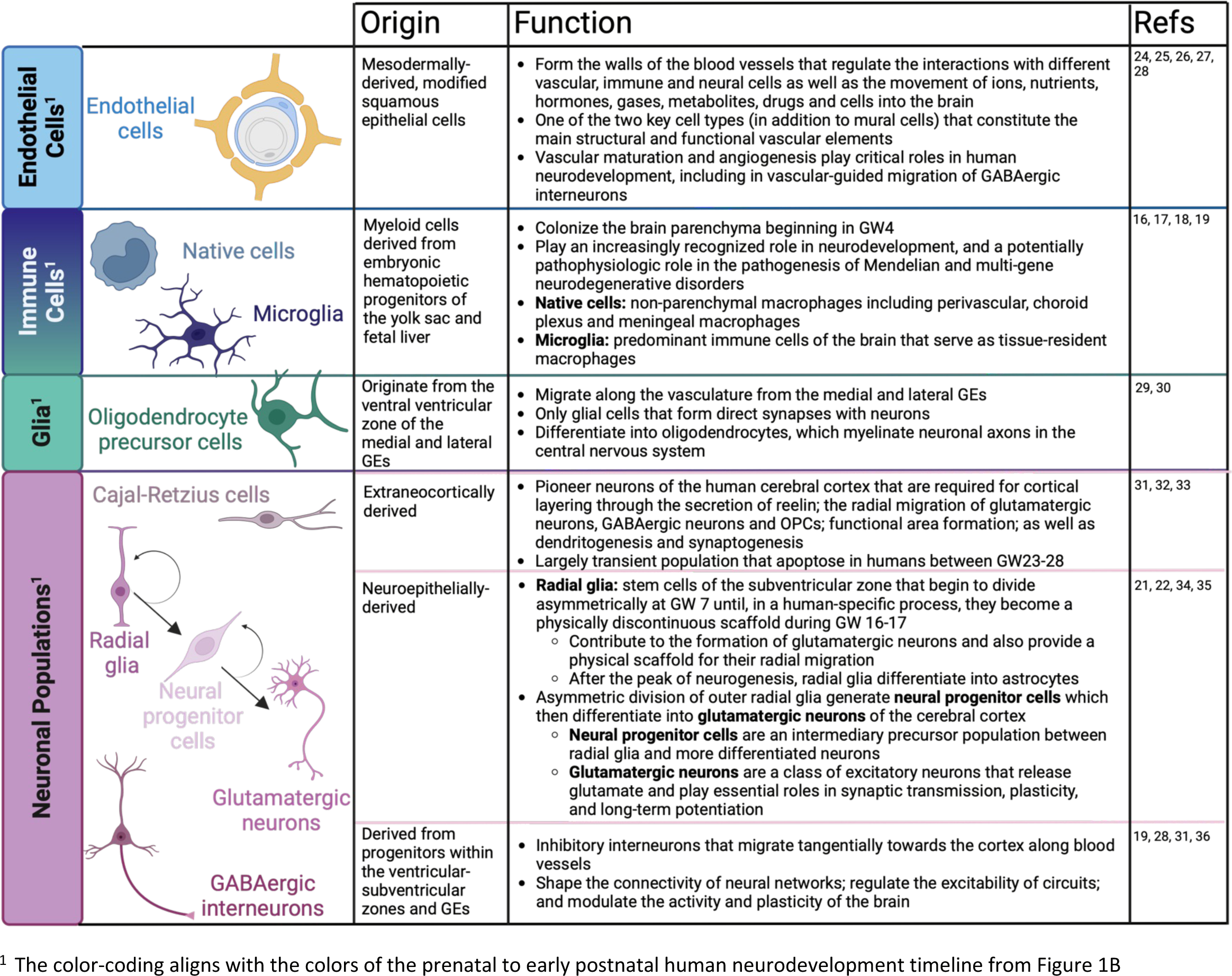
Primer on transcriptomically-delineated cell types in Bhaduri et al dataset.

There are thousands of unique cell types in the human brain. However, identifying all such cell types using single-cell transcriptomic data remains a difficult task. Single cell RNA-sequencing has a high degree of uncertainty, likely due to sequence-specific RNA degradation and random sampling of lowly expressed genes. This results in many expression values being “zero”^23^. Further, identification of cell types relies on unsupervised clustering followed by time-consuming manual annotation^23^. Thus, rare cell types may be lumped into other clusters, cell types with variable expression patterns may be misidentified, and differentiating continuous cell types must be put into discrete categories. For example, at some point NPCs become neurons. This is a continuous process, but a line must be drawn between the cell types at a discrete point. Cell atlases detailing the expression profiles of different tissues and cell types will be a key resource for this part of single cell data analysis for all model organisms, but they are still a work in progress for human cell types.

The dataset harnessed in NeuroTri2-VISDOT profiles 10 brain structures and 6 neocortical areas. These data are resolved to 10 unique cell types by clustering each sample by cell type and sub-clustering to identify more granular cell types^14^. We provide a primer on those cell populations to facilitate interpretation of the visualized data using NeuroTri2-VISDOT (Table 1), but these cells have been reviewed in more detail elsewhere^16–19,21,22,24–35,36^. We provide NeuroTri2-VISDOT as a free web-based interface (https://kellyclark.shinyapps.io/NeuroTri2-VISDOT/) and as an open-access R Shiny application via GitHub (https://github.com/kellyclark1/VISDOT) to visualize the expression patterns of genes across cell types throughout the second trimester in the developing human brain.

In this paper, we employ NeuroTri2-VISDOT in several different settings to evaluate its utility for the translational research community that focuses on Mendelian NDDs. First, we interrogate the expression patterns of the causative genes linked to the subset of chromatinopathies associated with Rubinstein-Taybi Syndrome (RSTS) (OMIM #613684; 180849). We then apply NeuroTri2-VISDOT to the spectrum of syndromes categorized as Noonan Syndrome-like RASopathies, including Noonan Syndrome (NS) (OMIM #163950; #610733; #616559; #613224; #616564; #605275; #619745; #615355; #618624; #618499; #611553); NS with multiple lentigines (OMIM #151100); NS-like disorder with loose anagen hair (OMIM #607721, 617506); Cardiofaciocutaneous Syndrome (OMIM #115150, 615279, 615280); Costello Syndrome (OMIM #218040); CBL-related RASopathy (OMIM #613563); wide spectrum RASopathy (OMIM #615278, 609942); MAPK1-related RASopathy (OMIM # 619087); and newly described NS-like RASopathies caused by variants in CDC42 and YWHAZ^37,38^. Finally, we employ NeuroTri2-VISDOT to an emerging class of NDDs caused by germline variants in histone genes^6,39^. Using NeuroTri2-VISDOT, we demonstrate that single cell RNA sequencing data from the developing human brain can be used to motivate the evidence-based selection of relevant cell populations to explore the pathogenic mechanism underlying Mendelian NDDs. Additionally, we show how systematic visualization of the temporal gene expression profiles with NeuroTri2-VISDOT may help elucidate the neurobiology underlying the neuro-phenotypic heterogeneity of historically grouped Mendelian NDDs.

## Results

### Using NeuroTri2-VISDOT

The NeuroTri2-VISDOT web-based interface has been designed to maximize the utility of the Bhaduri et al dataset for translational researchers (Supplemental Video 1). In the search box, an investigator can input the gene names for as many genes as they are interested in comparing using the notation from Ensembl GRCh38 Release 110 (Supplemental Figure 1A). Multiple genes must be space-separated, and gene names are case-sensitive. Users can opt to select or deselect any combination of cell types they are interested in exploring. All 10 cell types will always be represented in dot plots, but investigators may wish to deselect a cell type that has a higher expression than the other cell types for the line plots of percent and average expression. For example, the gene *PAFAH1B1 (LIS1)* is associated with the structural Mendelian NDD classical lissencephaly, a disorder primarily associated with cytoskeletal reorganization during neuron migration^40^. A large percent of second trimester Cajal-Retzius cells (∼80%) express *PAFAH1B1* (Supplemental Figure 1B, top), which can obscure the percent of other cell types expressing the gene. By removing Cajal-Retzius cells from the line plots, an investigator could more closely interrogate the pattern of *PAFAH1B1* expression in the nine other cell types (Supplemental Figure 1B, bottom).

To enhance customizability, investigators can also specify the scale of the plots, which enables the standardization of axes across plots. The two scales that can be customized are those for average expression scaled and percent expression. Average expression scaled is the normalized expression values for that gene in that cell type, scaled to a mean of zero. Percent expression is the percentage of a particular cell type with at least one read mapping to the gene of interest^41^. Because genes can be expressed at a low level but still have large impacts on cell function and identity, we chose to focus on percent of cells expressing genes of interest rather than average expression of the gene in each cell in subsequent analyses but the ability to visualize both dimensions of expression is retained in NeuroTri2-VISDOT. Investigators are also able to download not only the individual plots but also the data visualized in the plots if they are interested in performing additional analyses.

### Applying NeuroTri2-VISDOT to established NDD families

#### Chromatinopathies: A focus on Rubinstein-Taybi Syndrome (RSTS)

Chromatinopathies are caused by germline variants in genes that encode for epigenetic machinery: the readers, writers, erasers, and remodelers of the epigenome^42^. This group of Mendelian disorders is highly variable at both the genotypic and phenotypic level. Here, we focus on *CREBBP* and *EP300*, genes which encode histone acetyltransferases in which germline variants are causative for RSTS. Post-natal therapeutic intervention with histone deacetylase inhibitors rescues the markers of neurological dysfunction in mouse models of RSTS, emphasizing the value of more deeply understanding the pathogenesis of this disorder to better understand all chromatinopathies^42^. RSTS is a particularly elegant example of locus heterogeneity underlying phenotypic variability. Individuals who harbor variants in *EP300* tend to have less severe clinical features compared to those with *CREBBP* variants^43–45^. For example, intellectual disability is typically more severe in individuals with causative *CREBBP* variants, as is autism or autistic behaviors and epilepsy^43–46^.

This phenotypic severity seems to correlate with the second trimester single cell expression signatures (Figure 2A). Both genes show a peak in expression at GW18 in endothelial cells, reaching a percent expression of about 35% (Figure 2A, right, unfilled boxes). Interestingly, *CREBBP* is expressed in ∼48% of Cajal-Retzius cells while *EP300* is only expressed in about 27% of this cell type. This difference in percent expression, specifically in Cajal-Retzius cells, may contribute to the more severe neurological phenotypes of individuals with *CREBBP-*related RSTS compared to individuals with *EP300*-related RSTS. Notably, neuroimaging findings in individuals with RSTS include partial/total agenesis or hypodysgenesis of the corpus callosum, and straight gyrus hypoplasia have also been observed, though the genic contributions have been incompletely delineated^47^. The transcript patterns coupled with the neuroradiologic phenotypes suggest that taking a gene- and cell type-informed approach to interrogate the pathogenic mechanism underlying RSTS may be an appropriate choice to disentangle clinical heterogeneity.

**Figure 2.**
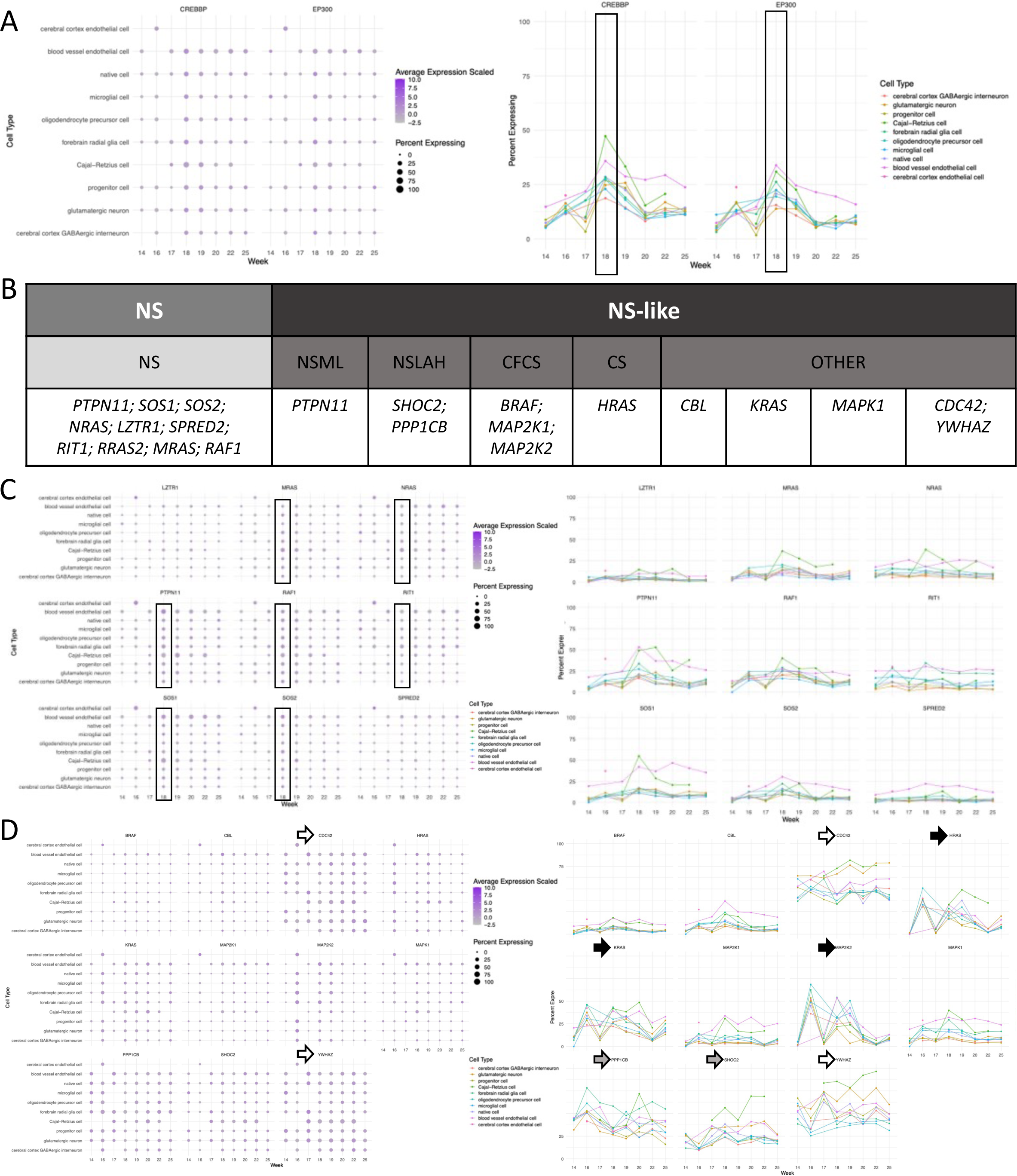
NeuroTri2-VISDOT captures genotypic and phenotypic heterogeneity of established NDDs. (A,C-D) Dot plots (left) – the average expression scaled reflects the normalized expression value for a gene, scaled to zero, and depicted by color gradient; the percent expression reflects the percentage of a particular cell type with at least one read mapping to the gene of interest, depicted by size gradient. Line plots (right) – percent expression by cell type. A) Dot plots (left) and percent expression line plots (right) for *CREBBP* and *EP300* across 10 neural cell types and 8 second-trimester time points. Unfilled boxes highlight GW18, emphasized in text. B) Table of genes associated with NS (left) and NS-like RASopathies (right). C) Dot plots (left) and percent expression line plots (right) for NS genes. Unfilled boxes highlight GW18, emphasized in text. D) Dot plots (left) and percent expression line plots (right) for NS-like RASopathy genes. Unfilled arrows highlight plots for the recently identified RASopathy genes, *CDC42* and *YWHAZ.* Gray arrows highlight plots for *SHOC* and *PPP1CB*, the genes underlying NSLAH. Black arrows highlight plots for *HRAS, KRAS* and *MAP2K2.* Abbreviations: NDDs = neurodevelopmental disorders; NS = Noonan Syndrome; NSML = NS with multiple lentigines; NSLAH = NS-like disorder with loose anagen hair; CFCS = Cardiofaciocutaneous Syndrome; CS = Costello Syndrome; GW = gestational week.

#### RASopathies: A focus on Noonan Syndrome and Noonan Syndrome-like (NS/NS-like) RASopathies

RASopathies, like chromatinopathies, are a genotypically and phenotypically heterogenous group of clinically overlapping Mendelian NDDs caused by variants affecting members and modulators of the RAS-MAPK cascade. Here, we focus on NS and NS-like RASopathies because of their universal classification as multisystem developmental disorders that often includes the CNS, which is not the case for non-NS-like RASopathies^38^. The characteristic inclusion of a neurologic phenotype in NS and NS-like RASopthies makes this variable group of syndromes amenable to interrogation with this neurodevelopmental dataset.

In concordance with previously published work, we define NS-associated genes as *PTPN11, SOS1, SOS2, NRAS, LZTR1, SPRED2, RIT1, RRAS2, MRAS* and *RAF1*^37,38^ (Figure 2B, left). Of the NS-like RASopathies, we define the associated genes as: *PTPN11* associated with NS with multiple lentigines; *SHOC2* and *PPP1CB* associated with NS-like disorder with loose anagen hair*; BRAF, MAP2K1* and *MAP2K2* associated with cardiofaciocutaneous syndrome; *HRAS* associated with Costello Syndrome; and *CBL*, *KRAS*, *MAPK1*, *CDC42*, and YWHAZ associated with other RASopathies^37,38^ (Figure 2B, right).

When we plot all NS/NS-like RASopathies together, it is challenging to identify any discernable patterns (Supplemental Figure 2). However, when we stratify based on the association of genes with NS compared to NS-like RASopathies, trends begin to emerge (Figure 2C, 2D). Nine of the 10 NS genes could be plotted (*RRAS2* was not able to be queried). Seven of these 9 genes have a peak at GW18 that transcends cell type (Figure 2C, left, unfilled boxes). Additionally, these same genes exhibit an upward trend in expression in both endothelial cells and Cajal-Retzius cells, which is consistent with what we observe when looking at the percent expression line plots (Figure 2C, right, pink and green). Taken together, there is a fairly consistent expression signature shared by genes that harbor causative variants in NS.

Compared to the NS genes, we observe heterogeneity in the second trimester brain expression of the NS-like RASopathy genes (Figure 2D). The first observation is that both the dot plots and the line plots for the newly described NS-like RASopathies associated with *CDC42* and *YWHAZ* do not follow the same trends as other NS-like RASopathy causative genes (Figure 2D, unfilled arrows). In general, the average and percent expression of these two genes throughout the second trimester is consistently greater than what we observe for any other gene queried in this NS-like RASopathy analysis (Figure 2D). The expression profiles visualized through the percent expression line plots is distinct in comparison to the other NS/NS-like RASopathy genes (Figure 2D, right, unfilled arrows). It has been previously noted that the assignment of the conditions caused by variants in these genes to the RASopathy family is under debate^38^. These data, along with other disparate features of *CDC42* and *YWHAZ-* related disorders, may support the reevaluation of the assignment of these conditions to the RASopathy family.

The value of looking at both the dot plots and the percent expression line plots is exemplified when probing the genes that underly NS-like disorder with loose anagen hair: *SHOC2* and *PPP1CB*. The dot plots for these genes look similar to each other, but distinct from other NS-like RASopathy genes (Figure 2D, left). However, the percent expression plots of these genes in different cell types across the second trimester are profoundly different. *SHOC2* has a bimodal peak in percent expression in Cajal-Retzius cells at GW18 and GW20 (Figure 2D, right, gray arrows). Conversely, *PPP1CB* has a single peak in percent expression at GW20 in Cajal-Retzius cells, as well as a peak in expression at GW16 in forebrain RG cells. As with the case of *CREBBP* or *EP300-*derived RSTS, these data may support the use of different model systems, for example Cajal-Retzius cells and/or forebrain RG, to study NS-like disorder with loose anagen hair driven by causative variants in different genes. Investigating pathogenic mechanisms through a bifurcated approach that centers on multiple cell types could enable the identification of a shared, downstream perturbation that could be targeted through therapeutic intervention.

Strikingly, the peak in expression at GW16 in forebrain RG cells that we observe in *PPP1CB*-related NS-like disorder with loose anagen hair is also present in percent expression plots for *MAP2K2*-related cardiofaciocutaneous syndrome, *HRAS*-related Costello Syndrome and *KRAS-*related wide spectrum RASopathy (Figure 2D, right, black arrows). These bimodal peaks in expression transcend NS-like RASopathy delineations; gene specification subgroups defined by the ClinGen RASopathy Expert Panel, in which genes with similar function and/or structure have been grouped^48^ (Supplemental Figure 3); and genes that share a gain-of-function disease mechanism (Supplemental Figure 4). Analysis of these expression signatures may prove useful for geneticists when making classification decisions about whether multiple syndromes should be grouped together in one family or into distinct entities.

### Applying NeuroTri2-VISDOT to an emerging class of NDDs

Germline variants in genes encoding histones are an emerging class of Mendelian NDDs^39^, which have recently been classified by OMIM. HIST1H1E syndrome/Rahman Syndrome (OMIM #617537) is caused by germline variants in the gene *HIST1H1E/H1-4* which encodes the histone H1 linker protein. Bryant-Li-Bhoj Syndrome (OMIM #619721, #619720) is caused by germline variants in *H3-3A* and *H3-3B*, the genes that encode histone H3.3. Tessadori-Bicknell-van Haaften NDD (OMIM #619758, #619759, #619950, #619551) is caused by germline variants in the genes *H4C3*, *H4C11*, *H4C5*, and *H4C9*, the genes that encode H4. Additional histone-encoding genes, including *MACROH2A1*, *H2AZ1*, *MACROH2A2*, *H2AZ2*, *H2AX*, and *H1-0*, are predicted to be putative disease candidates^6^.

The genes linked to HIST1H1E syndrome and Tessadori-Bicknell-van Haaften NDD are classified as replication-coupled (RC) histones, which are associated with unique structural and functional features. RC histone genes encode the only known cellular mRNA transcripts that are not poly-adenylated^6^. This lack of a polyA tail on RC histone transcripts renders them undetectable in the most common type of library preparation method used for RNA-sequencing. This means that in most publicly available datasets, including the one employed here, we are unable to explore their expression. Nonetheless, we are able to interrogate the expression of replication-independent (RI) histones that do have polyA tails. Based on prior work, we can stratify these RI histones into known NDD-causing, predicted NDD-causing, and non-disease-causing groups^6^ (Figure 3A).

**Figure 3.**
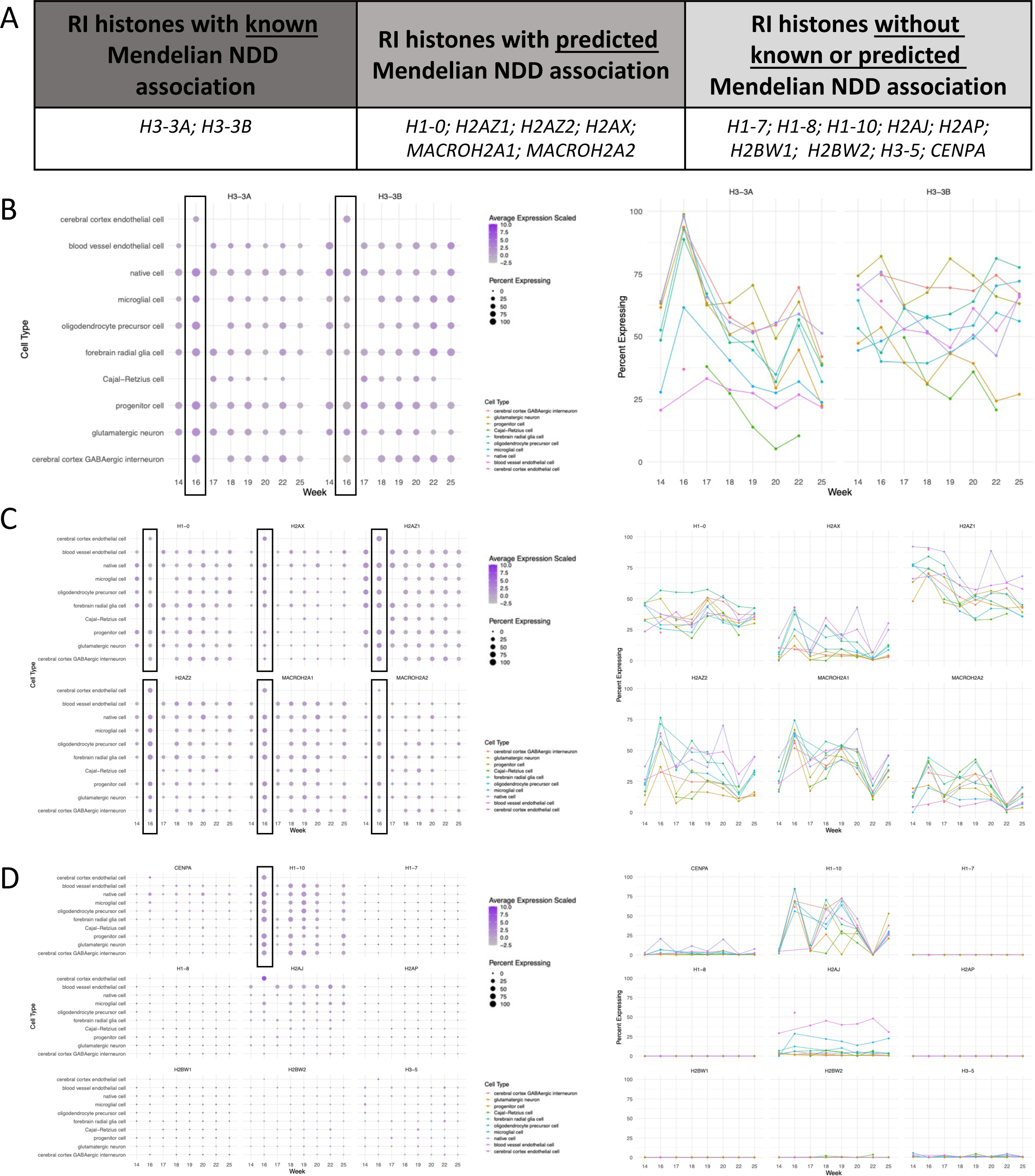
Novel application of NeuroTri2-VISDOT to histone-associated NDDs. A) Table of RI histone genes delineating which are known or predicted to be associated with NDDs. (B-D) Dot plots (left) – the average expression scaled reflects the normalized expression value for a gene, scaled to zero, and depicted by color gradient; the percent expression reflects the percentage of a particular cell type with at least one read mapping to the gene of interest, depicted by size gradient. Line plots (right) – percent expression by cell type. Unfilled boxes highlight GW16, emphasized in text. B) Dot plots (left) and percent expression line plots (right) for RI histone genes with known Mendelian NDD association. C) Dot plots (left) and percent expression plots (right) for RI histone genes with predicted Mendelian NDD association. D) Dot plots (left) and percent expression plots (right) for RI histone genes without known or predicted Mendelian NDD association. Abbreviations: NDDs = neurodevelopmental disorders; RI = replication-independent; GW = gestational week.

When we plot the expression of these RI histones, all the genes known or predicted to be associated with Mendelian NDDs show increases in expression across the second trimester, compared to genes not known or predicted to be associated with NDDs, with the exception of *H1-10* (Figures 3B-D).

Additionally, these genes seem to have a peak in expression at GW16 irrespective of cell type (Figure 3B-D, unfilled boxes). Interestingly, this is the window during which RG become physically discontinuous^21,22^. This expression pattern suggests an importance for RI histones at this very early point in brain development, and that RG would be an effective model to study the phenotypes associated with these variants. Neuroimaging results from individuals with HIST1H1E Syndrome, Bryant-Li-Bhoj Syndrome and Tessadori-Bicknell-van Haaften NDD do not show salient dysregulation of neural migration or cortical development. Thus, the role of RG in the neuropathology of these syndromes has not been explored. Through the visualization of these data using NeuroTri2-VISDOT, we identify a novel terminal lineage to which hiPSCs can be differentiated for subsequent functional work. More broadly, this demonstrates the utility of single cell expression signatures in development for identifying and supporting NDD candidate genes.

## Discussion

Clinical genetics is rooted in a rich history of clinical phenotyping that long predates the field’s ability to perform diagnostic genetic testing. In some cases, this has led to phenotypically similar syndromes being grouped together under large umbrella characterizations, such as NDDs or leukodystrophies, that may not reflect the distinct genetic processes contributing to the pathophysiology^1^. Now, the field of translational genetics is reckoning with this same question: to lump disorders together or split them into functional groups^49^. With NeuroTri2-VISDOT, we introduce a tool that may prove a valuable resource in this pursuit, with utility demonstrated for both established and emerging NDDs.

Single-cell RNA sequencing is an incredibly powerful method for exploring tissue heterogeneity and gene expression across cell types. However, generating a single-cell dataset is expensive and availability of human tissues, especially human fetal tissue, is limited. Further, data analysis requires computational skills and high-performance computing due to the size and complexity of the multidimensional data produced in these experiments. NeuroTri2-VISDOT seeks to address the computational barriers to use of this valuable tool, particularly the size of single-cell data and the computational skills needed to explore it. First, the complete dataset utilized here is available as a 50GB file, which is too large for many researchers to store and use on their own computers necessitating the use of high-performance computing. NeuroTri2-VISDOT stores a summary of this dataset online, with average scaled expression and percent expression for each gene by cell type rather than read counts for each cell. No downloads are necessary to use the web app, and the raw data for genes of interest can be downloaded as a small file of comma-separated values, reducing the need to download the entire dataset. Second, analysis of single cell data using the R package, Seurat, requires many researchers to dedicate significant time to learning how to navigate R and Seurat objects. NeuroTri2-VISDOT reduces the need for coding expertise by allowing exploration of single cell data with a user-friendly interface that requires only a gene or list of genes as input.

We recognize some limitations associated with our approach. First, this dataset was generated from single-cell transcriptomics performed on microdissected regions of the brain that included 10 major forebrain, midbrain and hindbrain regions in addition to 6 neocortical areas, meaning that only cell types identified in the neocortical samples were powered to be reported in the final dataset^14^. Thus, some cellular populations important to neurodevelopment, such as cells of the cerebellum, are not represented. Astrocytes, which differentiate from RG after the peak of neurogenesis at the end of the second trimester, are also not represented. These cells play an important homeostatic role in the brain parenchyma and form the neural component of the blood brain barrier. However, there are thousands of unique cell types in the human brain, and it is not feasible to capture them all. Further, while these data provide a powerful window into neurodevelopment by quantifying gene expression in the second trimester, earlier and later stages of prenatal brain development are not captured here. Nonetheless, NeuroTri2-VISDOT is intended not as an atlas of all cells in all stages of the developing brain, but rather as a tool to efficiently leverage this powerful dataset in translational genetic research.

Another limitation of this analysis is the interdependence and interconnectivity of different cell types during neurodevelopment. For instance, GABAergic interneurons and oligodendrocyte precursor cells require the endothelial-lined vasculature to appropriately migrate. Additionally, NPCs not only arise from but also migrate along RG before eventually differentiating into glutamatergic cortical neurons. This complexity, in part, motivates some investigators to opt for model organisms, such as mice or zebrafish as opposed to hiPSCs. The expression visualization made possible by NeuroTri2-VISDOT captures this complexity, but also enables investigators to make evidence-based decisions about model selection. For instance, if RG are nominated as a compelling model to interrogate, it is crucial to consider that this cell type undergoes the process of becoming a physically discontinuous scaffold during the middle of the second trimester, which may support the use of hiPSCs over mice and zebrafish in specific situations. These data may also indicate that different cell populations are worth interrogating based on which gene harbors variants driving the phenotype, as in the case of *CREBBP*-versus *EP300*-driven RSTS.

A biological consideration before employing these data is whether the RNA transcript is the appropriate read out for a given gene of interest. Transcript and protein levels are discordant throughout neurodevelopment, most prominently in post-natal life^50^. When exploring post-natal neurodevelopment, protein or phosphorylated protein may be a more precise metric to inform model selection. In cases where transcript level expression is an appropriate biologic read-out, such as in pre-natal neurodevelopment, we propose that there are several ways in which NeuroTri2-VISDOT could be applied in the future (Figure 4). At the level of an individual translational research lab, we envision NeuroTri2-VISDOT as a tool to enable investigators to systemically inform the selection of their model systems. We see this approach being adapted by bioinformaticians to create similar tools for the increasingly available datasets designed to ask similarly targeted questions. For example, a similar tool to visualize neural gene expression across cell types across the lifespan could be employed by biobanks to identify and cluster gene expression signatures of putative disease candidate genes.

**Figure 4.**
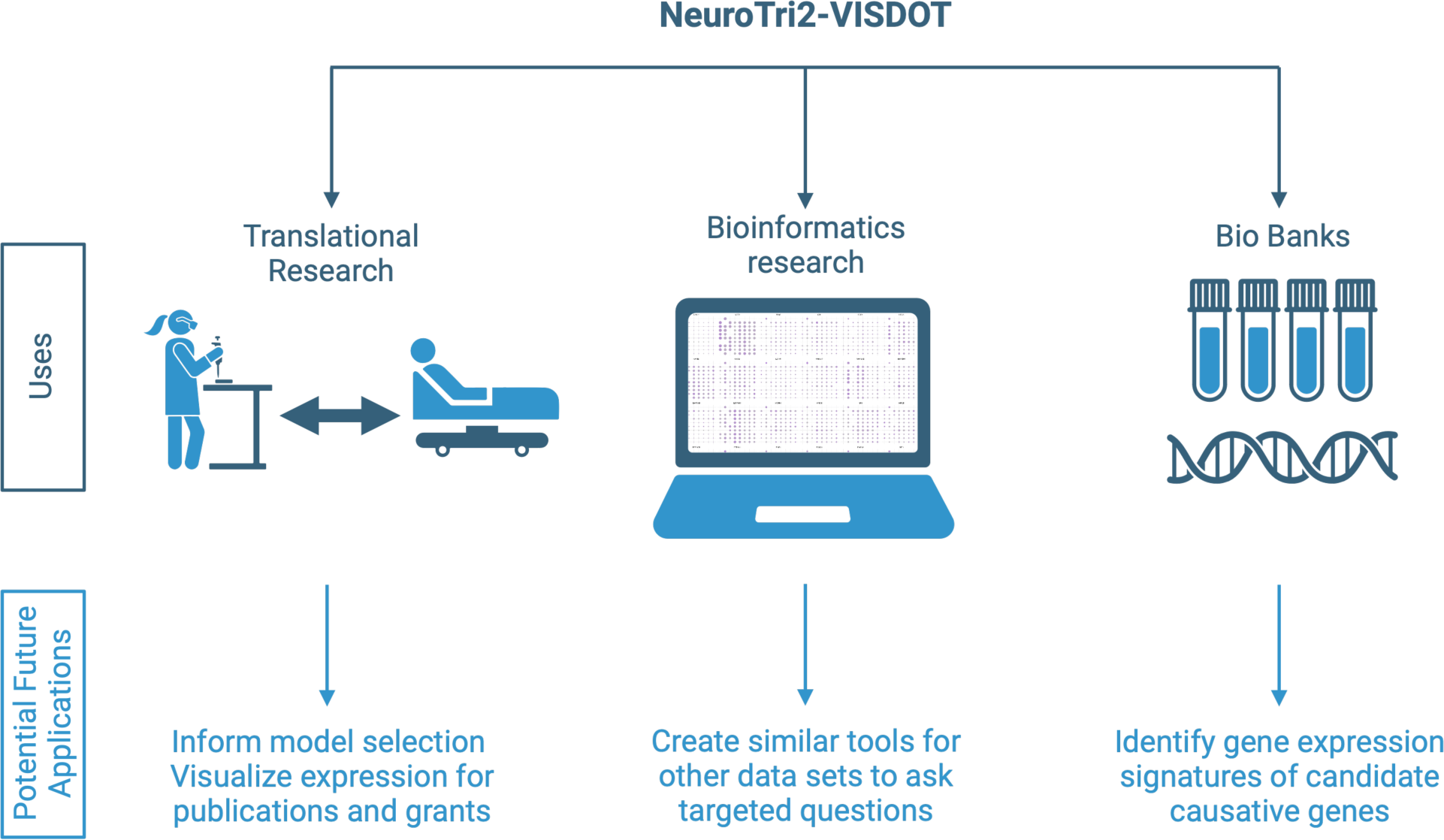
Uses and future applications of NeuroTri2-VISDOT. NeuroTri2-VISDOT can be used, as demonstrated, at the level of a translational genetic research lab to inform model selection. More broadly, similar tools can be built to address the same questions in other tissues and at other developmental stages using different data sets. Expression signatures can also be compared across multiple genes in BioBanks to nominate candidate causative genes.

In summary, the numerous applications of NeuroTri2-VISDOT, including model selection and interrogation of gene signatures across time and cell types, allow for translational genetics researchers to harness the power of a valuable single-cell second trimester brain dataset without requiring programming experience or access to high performance computing resources. The approach employed to generate and evaluate NeuroTri2-VISDOT also provides a framework for developing tools for similar datasets that will be beneficial to the wider translational research community.

## Methods

### Data extraction

The Seurat object for the data generated by Bhaduri et al. is publicly available^14,41^. This object was split into separate objects for each cell type, and plots were generated for genes of interest for each cell type using the Seurat DotPlot function. The data in these dot plots were combined to create a large table containing each gene with the percent expression and average expression scaled at each time point in each cell type. This data is available to download from the NeuroTri2-VISDOT web app. Additionally, the code used to generate NeuroTri2-VISDOT can be found at https://github.com/kellyclark1/VISDOT.

### NeuroTri2-VISDOT Web app development

The NeuroTri2-VISDOT web app was written in R v4.2.2^51^ using the R package shiny v1.7.3^52^ and is available at https://kellyclark.shinyapps.io/NeuroTri2-VISDOT/. Plots displayed on the web interface were designed using ggplot2 v3.4.0^53^.

The web app requires as input at least one human Ensembl GRCh38 Release 110 gene name and will accept as many gene names as a user would like to compare. Users may also select cell types of interest for line plots using the checkboxes in the left panel. The web app allows custom scaling for plots, which should be used with caution. The intent of custom scaling is to allow multiple genes to be plotted separately but at the same scale. However, setting limits that do not include the data will exclude data points outside of those limits.

## Supporting information

Supplemental Video 1 Transcript

## Data Availability

Data and code used to generate NeuroTri2-VISDOT can be found at https://github.com/kellyclark1/VISDOT.

## Acknowledgements

We would like to thank Dr. Christopher Sifuentes of the Chan Zuckerberg Initiative not only for his invaluable insights pertaining to data wrangling visualization but also his tireless support of trainees.

## Author Contributions

CRediT has 14 categories:

1. Conceptualization – KJC, EEL, EMG, TTN, EBJ
2. Data curation – EEL, KJC, EMG
3. Formal analysis – KJC, EEL, EMG
4. Funding acquisition – RAN, EJB
5. Investigation – KJC, EEL, EMG
6. Methodology – KJC, EEL, EMG, TTN, EJB
7. Project administration – TTN, EJB
8. Resources – RAN, EJB
9. Software - KJC
10. Supervision – TTN, EJB
11. Validation – KJC, EEL, EMG, AKS, DLC, ELD, RAN, TTN, EJB
12. Visualization – KJC, EEL
13. Writing-original draft – EEL, KJC
14. Writing-review & editing – EEL, KJC, EMG, AKS, DLC, ELD, RAN, TTN, EJB

## Ethics Declaration

This study does not involve human subjects or live invertebrate and/or higher invertebrates.

