## Supplemental Figures 1-4 for "NeuroTri2-VISDOT: An open-access tool to harness the power of second trimester human single cell data to inform models of Mendelian neurodevelopmental disorders"

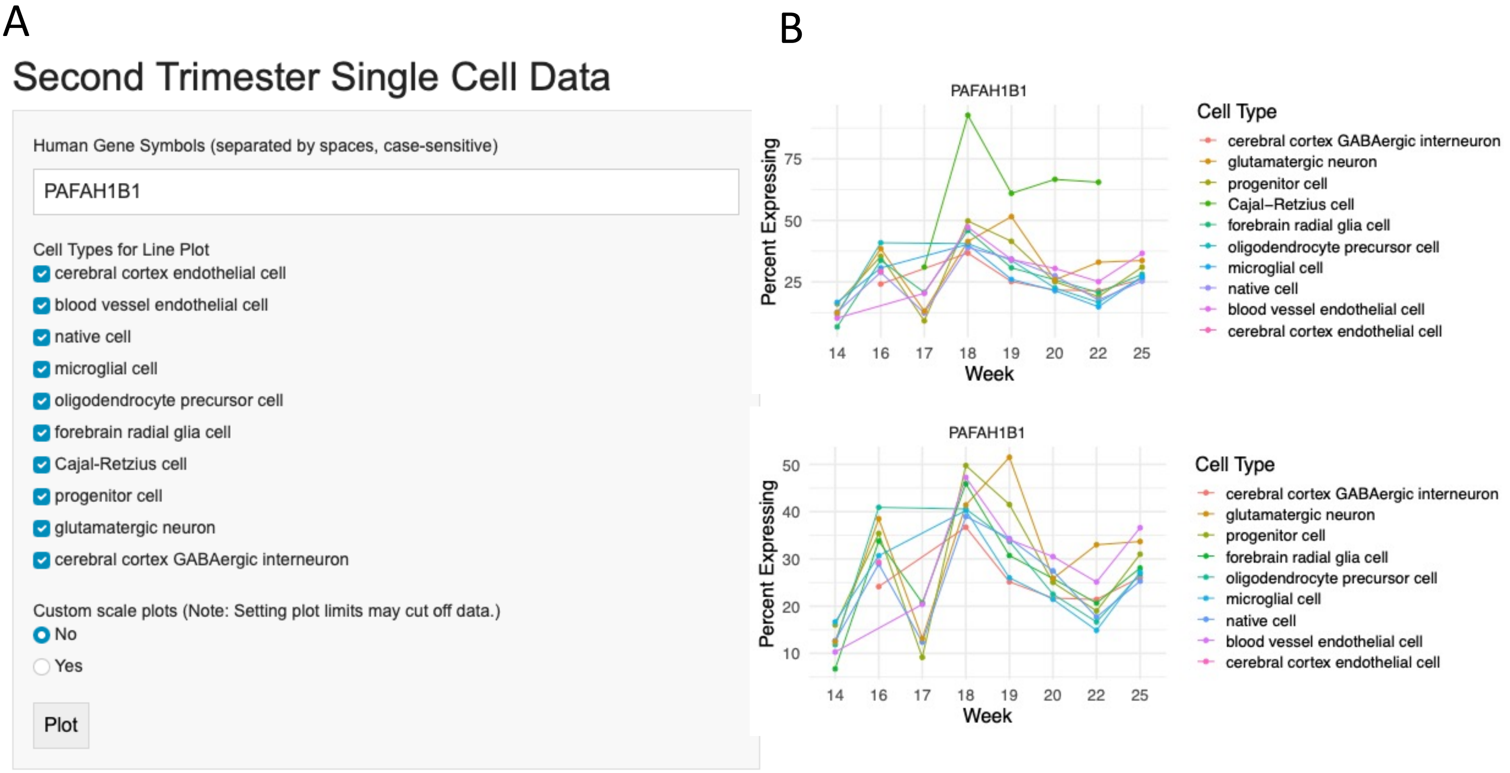


**Supplemental Figure 1 NeuroTri2-VISDOT interface and usage.** A) Search feature requiring input of capitalized, space-separated gene names using notation from Ensembl GRCh38 Release 110. Customizable line plot features include selection/deselection of cell types and scaling. B) Percent expression line plots showing cell type selection/deselection with *PAFAH1B1*. Top – all cell types selected. Bottom – Cajal-Retzius cells deselected allowing for the expansion of the y-axis so the expression of more lowly-expressed cell types can be more clearly visualized.

A

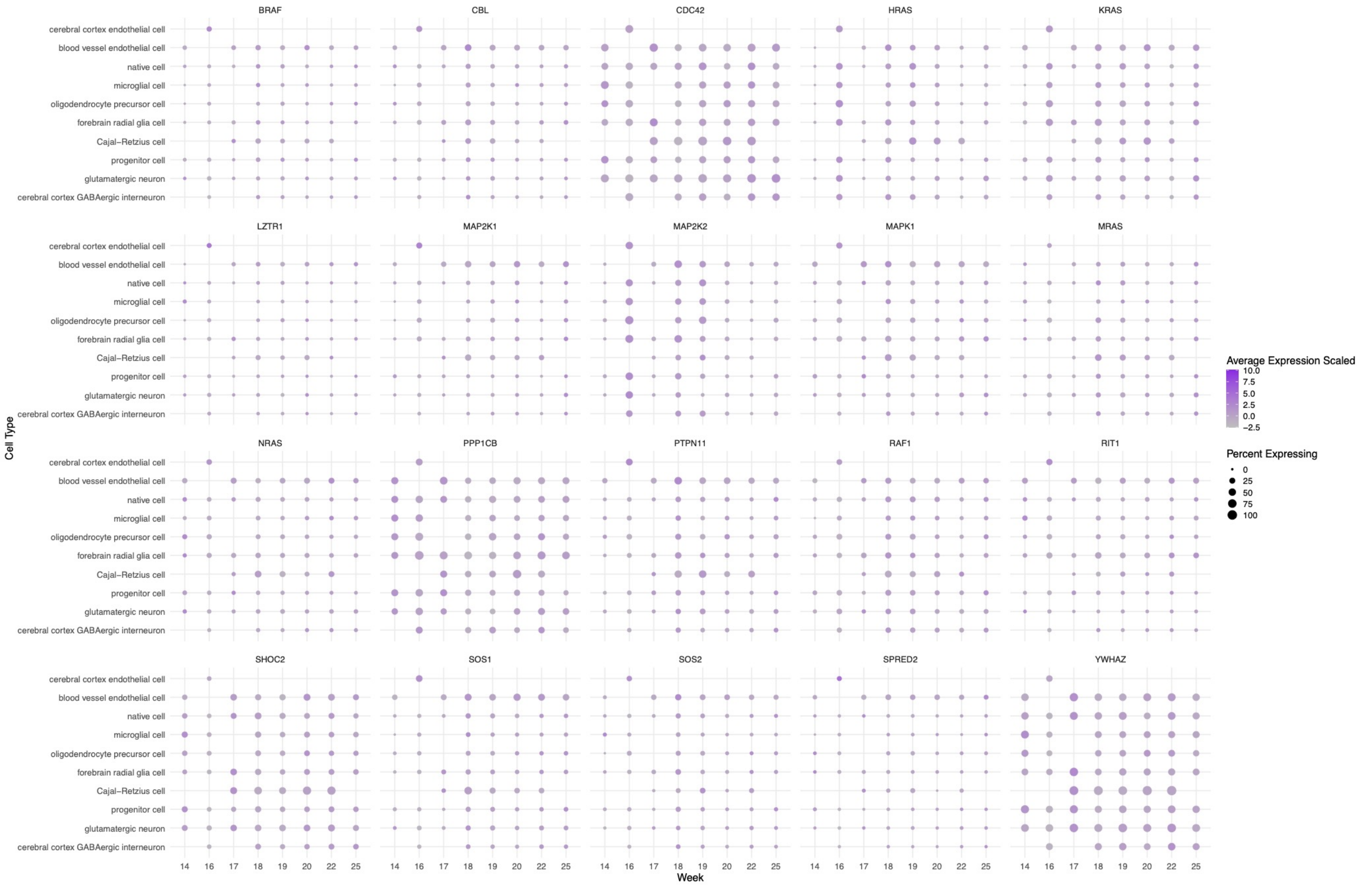


B

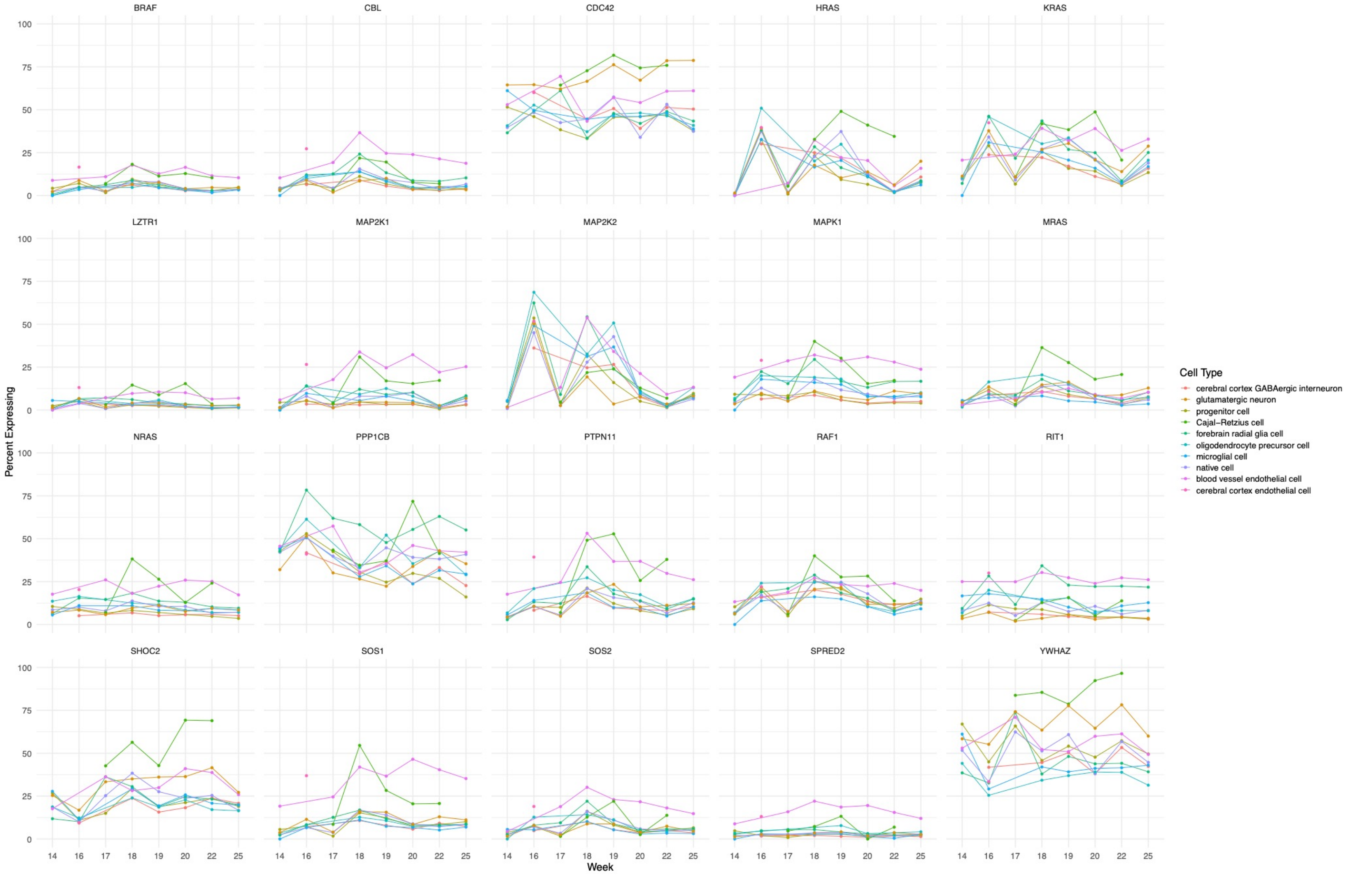


**Supplemental Figure 2 Unstratified visualization of all NS and NS-like RASopathy genes.** Without stratification, expression patterns are not discernable. A) Dot plots of all NS and NS-like RASopathy genes. The average expression scaled reflects the normalized expression value for a gene, scaled to zero, and depicted by color gradient; the percent expression reflects the percentage of a particular cell type with at least one read mapping to the gene of interest, depicted by size gradient. B) Percent expression line plots of all NS and NS-like RASopathy genes. Percent expression by cell type by time.

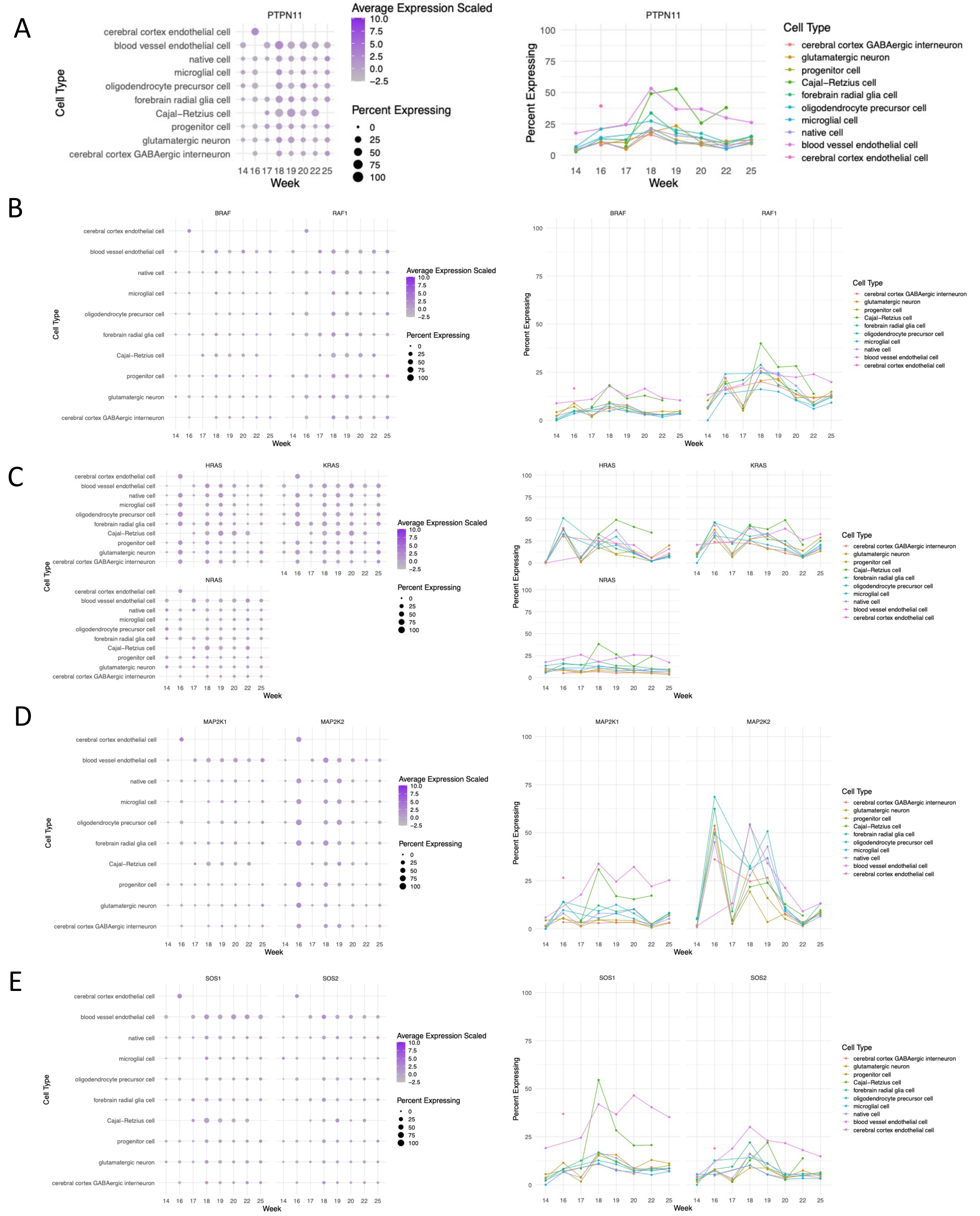


**Supplemental Figure 3 Gene specific subgroups based on shared function and/or structure identified by ClinGen's RASopathy Expert Panel panel consensus.** Bimodal gene expression peak in RG is not a defining characteristic of any one of these subgroups yet we see that signature shared by different genes in each group. Five subgroups: A) *PTPN11*, B) *BRAF/RAF1*, C) *HRAS/KRAS/NRAS*, D) *MAP2K1/MAP2K2I,* E) *SOS1/SOS2.* Dot plots (left) – the average expression scaled reflects the normalized expression value for a gene, scaled to zero, and depicted by color gradient; the percent expression reflects the percentage of a particular cell type with at least one read mapping to the gene of interest, depicted by size gradient. Line plots (right) – percent expression by cell type.

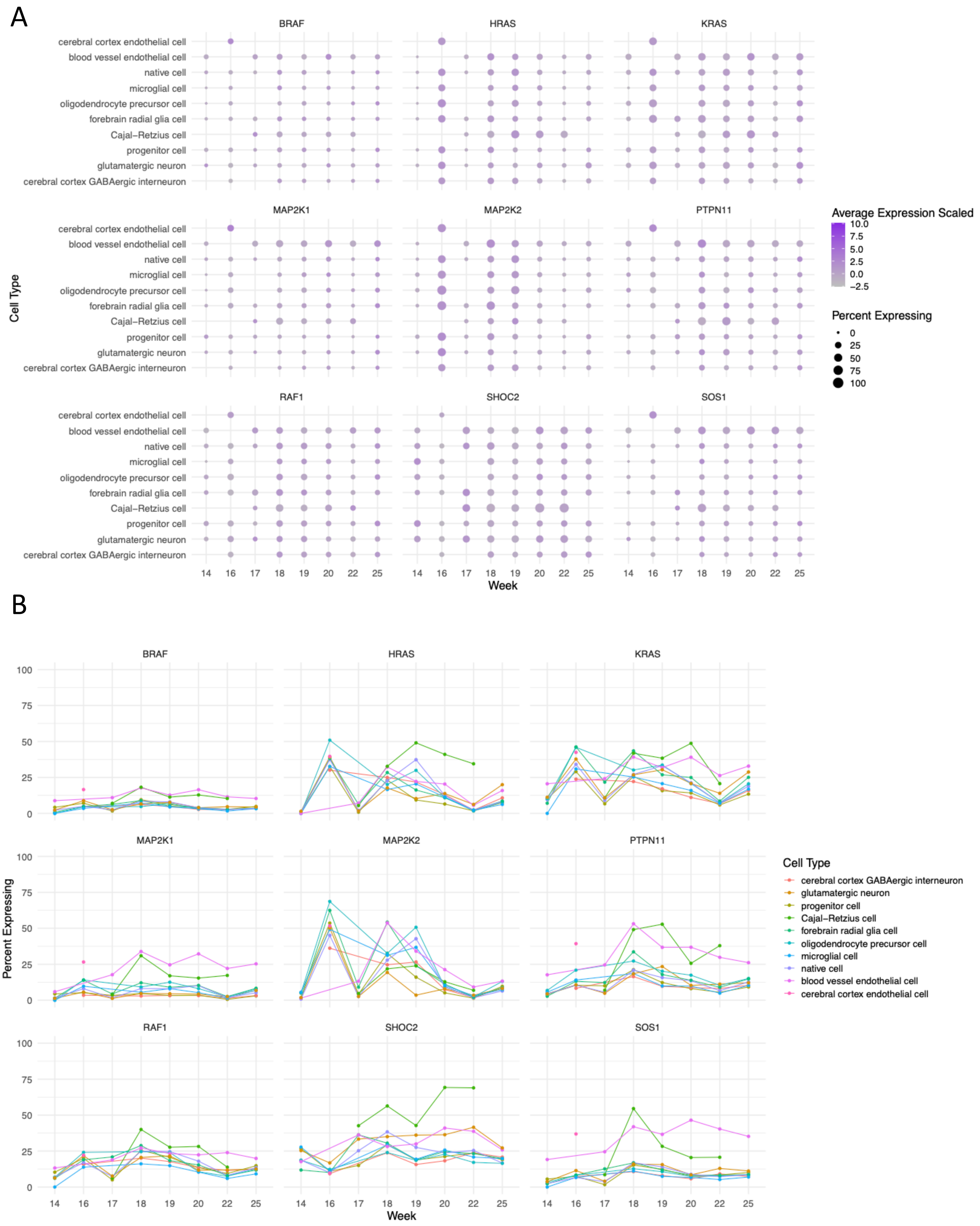


**Supplemental Figure 4 NS and NS-like genes with gain-of-function disease mechanism.** Bimodal gene expression peak in RG present in some but not all NS and NS-like genes with gain-of-function disease mechanism. A) Dot plots - the average expression scaled reflects the normalized expression value for a gene, scaled to zero, and depicted by color gradient; the percent expression reflects the percentage of a particular cell type with at least one read mapping to the gene of interest, depicted by size gradient. B) Percent expression line plots - percent expression by cell type.
