## Supplemental Video 1 Transcript for "NeuroTri2-VISDOT: An open-access tool to harness the power of second trimester human single cell data to inform models of Mendelian neurodevelopmental disorders"

This is the NeuroTri2-VISDOT web interface. To plot a gene, type the HGNC gene symbol into the text box and click plot. Here we have the dot plot, the percent expressing line plot, and the scaled average expression line plot. If you want to view only specific cell types in the line plots, you can check and uncheck the cell type boxes. For example, I can exclude Cajal-Retzius cells, and when I click plot again, the line plots will rescale. If I want to change the plot scale for both dot and line plots, I can select yes here, then input the numbers I'd like to change. If you input limits that do not include all the data points, data outside those limits will be excluded. You can also plot multiple genes, by inputting their gene symbols separated by spaces. If you want to export plots, these download buttons will download the plots as pdf files. If you want to export the data for your genes of interest, the data button can be used to download the data as a csv file. The about button contains citation information.
